# Plant metabolites modulate animal social networks and lifespan

**DOI:** 10.1101/2023.12.20.572488

**Authors:** Pragya Singh, Leon Brueggemann, Steven Janz, Yasmina Saidi, Gaurav Baruah, Caroline Müller

## Abstract

Social interactions influence disease spread, information flow, and resource allocation across species, yet heterogeneity in social interaction frequency and its fitness consequences remain poorly understood. Additionally, animals can utilize plant metabolites for purposes beyond nutrition, but whether that shapes social networks is unclear. Here, we investigated how non-nutritive plant metabolites impact social interactions and the lifespan of the turnip sawfly, *Athalia rosae*. Adult sawflies acquire neo-clerodane diterpenoids (’clerodanoids’) from non-food plants, showing intraspecific variation in natural populations and laboratory-reared individuals. Clerodanoids can also be transferred between conspecifics, leading to increased agonistic social interactions. Network analysis indicated increased social interactions in sawfly groups where some or all individuals had prior access to clerodanoids. Social interaction frequency varied with clerodanoid status, with fitness costs including reduced lifespan resulting from increased interactions. Our findings highlight the role of intraspecific variation in the acquisition of non-nutritional plant metabolites in shaping social networks, with fitness implications on individual social niches.

## Introduction

Social interactions are widespread in animals and lead to the emergence of diverse social networks, from nominally to highly interactive (Frank 2007; Krause *et al*. 2015). Such interactions can range from mating to agonistic. A common aspect of networks constructed on social interactions, i.e. animal social networks, is skewness in the number of interactions individuals have (Fisher *et al*. 2019; Gartland *et al*. 2022; Krause *et al*. 2015). Some individuals have frequent social interactions while others have few. Consequently, this could have ecological ramifications. For instance, individuals with a higher frequency of interactions may overly affect disease transmission (Lloyd-Smith *et al*. 2005), which in turn may shape evolution of social systems (Udiani & Fefferman 2020). Furthermore, more or few social interactions may have associated benefits or costs for an individual (Koto *et al*. 2023; Ruan & Wu 2008; Snyder-Mackler *et al*. 2020). For example, increased agonistic social interactions can lead to injuries or even mortality (Archer 1988), although such costs are more widely documented in species that have teeth, claws or other armaments than in unarmed species. Even in unarmed species, there can be costs of frequent social interactions, such as depletion of energy reserves (Briffa & Sneddon 2007), such as reduced lipid or carbohydrate reserves.

Social interactions can depend on various biotic and abiotic factors (Candolin & Wong 2012; Fisher *et al*. 2021). One of these biotic factors may be plant-animal associations, but their role in explaining the observed skew in social interactions is only little explored. Studies have shown that several specialised plant metabolites are taken up by certain animal species independently of nutrition, and stored (sequestered), for example, for use in mating or as anti-predation defence (Beran & Petschenka 2022; Nishida 2014; Opitz & Müller 2009). This phenomenon is called pharmacophagy (Boppré 1984). Remarkably, such non-nutritive plant metabolites or their derivatives, can also be acquired from other animals, either conspecifics (Paul *et al*. 2021; Singh *et al*. 2022) or heterospecifics (Tea *et al*. 2021). The uptake of such non-nutritive plant metabolites can have direct effects on the behaviour of the individuals (Amano *et al*. 1999; Conner *et al*. 2000; Gonzalez *et al*. 1999). This individual-level variation in behaviour and how the individuals interact with each other can in turn lead to the emergence of complex patterns in animal social networks (Krause *et al*. 2015). Thus, these sequestered plant metabolites can impact social interactions, and consequently influence the fitness of the individual animals (Formica *et al*. 2012; Royle *et al*. 2012; Silk *et al*. 2003) and their individualised social niche (Kaiser *et al*. 2024; Trappes *et al*. 2022). Moreover, taking up plant metabolites may entail other costs, such as potential toxicity, if the substances are harmful (Kortbeek *et al*. 2019).

Interestingly, many animal species show intraspecific variation in the presence and quantity of sequestered plant metabolites (Liu *et al*. 2023; Mattila *et al*. 2021; Opitz & Müller 2009; Speed *et al*. 2012). Such variation can result from variation in the plant metabolite concentration per se, potential costs related to metabolite acquisition by the animal or factors such as age, sex or immunological status of the individuals (Dimarco & Fordyce 2017; Massad *et al*. 2011; Moore *et al*. 2014; Smilanich *et al*. 2009; Zvereva & Kozlov 2015). This raises the question of whether intraspecific variability in sequestered plant metabolites can influence social interactions, and if these metabolites, directly or through modified social interactions, impact fitness indicators such as lifespan. Additionally, the transfer of plant metabolites between individuals is likely to influence the social interactions of both the donor and the recipient. However, despite the known impact of sequestered plant metabolites on individual behaviour (Boppré 1984; Nishida 2014), our understanding of how these effects translate into broader implications for social interactions remains limited.

To target these open questions, we examined the effect of intraspecific variation in sequestered plant metabolites on social interactions and lifespan in the turnip sawfly, *Athalia rosae* (Hymenoptera: Tenthredinidae). The adults of *A. rosae* feed on floral nectar, usually from Apiaceae plants. In addition, they visit non-food plant species such as *Ajuga reptans* (Lamiaceae), on which they exhibit pharmacophagy by taking up specialised metabolites, neo-clerodane diterpenoids (hereafter called ‘clerodanoids’), which they further metabolise slightly (Brueggemann *et al*. 2023). These (metabolised) clerodanoids can serve as a resource for *A. rosae* adults in multiple ways (Paul *et al*. 2022). Access to clerodanoids increases their mating success (Amano *et al*. 1999) and also provides chemical defence against predators (Nishida & Fukami 1990; Singh *et al*. 2022). Moreover, individuals can obtain clerodanoids via agonistic interactions like nibbling on conspecifics that have had access to plant clerodanoids (Paul & Müller 2021; Singh *et al*. 2022). Such agonistic interactions can potentially be costly, as suggested by the transcriptional upregulation of metabolic pathways in sawflies that nibbled on conspecifics (Paul *et al*. 2021). Additionally, the uptake of clerodanoids may also directly pose fitness costs, such as a reduced lifespan (Zanchi *et al*. 2021). However, laboratory cultures of *A. rosae* are usually maintained without access to clerodanoids, indicating that clerodanoids are not essential for individual survival.

In this study, we initially assessed intraspecific variation in clerodanoid levels within natural *A. rosae* populations. Subsequently, we investigated how access to clerodanoids, either directly through *A. reptans* leaves or indirectly via conspecifics, influenced behavioral interactions in laboratory-reared individuals, and analysed their clerodanoid levels. Notably, intraspecific variation in clerodanoid content was observed in both wild-caught and laboratory-reared individuals. To further understand the impact of clerodanoid variation on social interactions, we established treatment groups with different levels of clerodanoid access, using network analysis to quantify individual and group social interaction metrics (Wey *et al*. 2008). Lastly, we explored the fitness consequences of clerodanoid acquisition by examining its effects on lifespan and metabolic profiles, i.e. lipid and carbohydrate content, employing treatment groups with varying clerodanoid levels. Through these analyses, we elucidated the ecological significance of clerodanoids in shaping social networks and individual fitness in *A. rosae* populations. We expected intraspecific variation in adult clerodanoid levels. Groups lacking clerodanoid access were expected to exhibit minimal social interactions, whereas asymmetric clerodanoid access would promote increased social interactions. Individuals without direct access were predicted to adjust social interaction frequency based on the clerodanoid status of conspecifics. Furthermore, increased social interactions for clerodanoid acquisition were expected to incur fitness costs, resulting in a reduced lifespan and a greater depletion of lipid and carbohydrate reserves.

## Materials and Methods

### Maintenance of study organism

Adults of *A. rosae* used in the experiments (other than wild-caught adults) were taken from a laboratory stock population that had been established from sawflies collected in and around Bielefeld, Germany, and annually supplemented with field-caught individuals (see Singh *et al*., 2023 for laboratory rearing details). The individuals were provided a 2% (v/v) honey-water solution on paper that was refreshed every alternate day. The laboratory population was maintained in multiple mesh cages (each 60 x 60 x 60 cm) at room temperature (15-25°C) with a 16:8 h light:dark cycle and approximately 60% relative humidity.

### Experiment 1. Chemical analysis for clerodanoid content of wild-caught sawflies

In total eighteen female and eight male adult *A. rosae* were collected in the wild at three locations (SI.1). For analysis of clerodanoid amounts, the insects were individually frozen, lyophilised, homogenised and extracted in ethyl acetate (LC-MS grade, VWR, Leuven, Belgium), using ultrasonication. After centrifugation, supernatants were dried and resolved in 100 µl of methanol with mefenemic acid (Sigma-Aldrich GmbH, Taufkirchen, Germany) as internal standard. Samples were filtered and analysed using an ultra-high performance liquid chromatograph (UHPLC; Dionex UltiMate 3000, Thermo Fisher Scientific, San José, CA, USA) equipped with a Kinetex XB-C18 column (1.7 µm, 150 × 2.1 mm, with guard column, Phenomenex) kept at 45 °C, coupled to a quadrupole time of flight mass spectrometer (QTOF-MS/MS; compact, Bruker Daltonics, Bremen, Germany) in negative electrospray ionisation mode, following Brueggemann *et al*. (2023; for details see Supplement).Samples were separated with a gradient from 0.1% formic acid (p.a., eluent additive for LC-MS, ∼98%, Sigma-Aldrich) in deionised water (eluent A) to 0.1% formic acid in acetonitrile (LC-MS grade, Fisher Scientific, Loughborough, UK; eluent B) at a flow rate of 0.5 ml min^-1^, starting eluent B at 2%, increasing to 30% within 20 min, and then to 75% within 9 min, followed by column cleaning and equilibration. The settings for the MS mode were capillary voltage 3000 V, end plate offset 500 V, nebuliser (N_2_) pressure 3 bar, dry gas (N_2_, 275 °C) flow 12 L min^-1^, quadrupole ion energy 4 eV and collision energy 7 eV. Line spectra were captured at 6 Hz, with a range of 50-1300 *m*/*z*. For recalibration of the *m*/*z* axis, a Na(HCOO)-solution was injected prior to each sample. Blanks without animal material were prepared in the same way and measured under the same conditions.

Mass axis recalibration and peak picking with the T-ReX 3D algorithm including spectral background subtraction were performed in Metaboscape 2021b (Bruker Daltonics), with an intensity threshold of 1000 counts and minimum peak length of 9 spectra. The allowed ion types for bucket generation were: [M-H]^-^, [M-H2O-H]^-^, [M+Cl]^-^, [M+HCOOH-H]^-^, [M+CH_3_COOH-H]^-^ and [2M-H]^-^. From each bucket only the feature with the highest intensity was used. We focussed on two metabolites which are likely taken up from *A. reptans* and metabolised by the sawflies (Brueggemann *et al*. 2023), putative clerodanoid 1 eluting at 16.3 min (*m/z* 529.229 [M+HCOOH-H]^-^) and putative clerodanoid 2 eluting at 18.2 min (*m/z* 527.214 [M+HCOOH-H]^-^), with the monoisotopic masses of 484.231 Da (proposed sum formula C_24_H_36_O_10_) and 482.214 Da (C_24_H_34_O_10_), respectively. Peak intensities of these target metabolites were related to the peak intensity of mefenemic acid for semi-quantitative analysis.

### Experiment 2. Effect of direct and indirect clerodanoid acquisition on social interactions

We investigated the effects of no clerodanoid access (C-), indirect clerodanoid access from a conspecific (C+ from conspecific), or direct clerodanoid access from a plant (C+ from plant) by females on social interactions with C- males. Note that ‘C+ from conspecific’ has been previously referred to as AC+ (Paul *et al*. 2021; Singh *et al*. 2022). For the ‘C+ from plant’ treatment, freshly eclosed females were provided individually an *A. reptans* leaf disc (∼1 cm^2^) for 48 h, while ‘C+ from conspecific’ treatment females were kept with a C+ conspecific (2-4 days old female that had previously contact to *A. reptans*) for 48 h. Males (6-10 days post-eclosion) were kept individually in Petri dishes for 48 h prior to the assays. The males were mated to a non-focal C- female (∼3 days old) 24 h prior to the experiment to avoid any differences arising due to first mating, as observed in other species (Torres-Vila & Jennions 2005). For the behavioural assays, one female was set up with one male in a clean Petri dish (5.5 cm diameter) and recorded (Sony HDR-CX410VE camcorder, AVCHD – 1920 × 1080 − 25 fps) for 25 minutes (16 replicates per treatment combination). One to two replicates of each treatment were set up simultaneously in a trial. An observer scored the number of occurrences of agonistic behaviour, including frontlimb battling, fighting and nibbling events (see Paul & Müller, 2021 for detailed behavioural descriptions), and mating behaviour (number of mating, latency until first mating) in each movie replicate using the software BORIS v7.13.9 (Friard & Gamba 2016), being blind to the treatment. After the behavioural assays, five females from each treatment were chosen randomly, frozen and analysed for their clerodanoid content as in experiment 1.

### Experiment 3. Effect of intraspecific variation in clerodanoids on social networks

We found intraspecific variation in clerodanoid content in both wild-caught and lab-reared sawflies (see Results). To test the effect of clerodanoid acquisition status on social networks, we set up three group types of two males and two females each, in which either no individual had prior access to clerodanoids (C-group, symmetric), only one female and one male had prior access to clerodanoids (Mixed group, asymmetric) or all individuals had prior access to clerodanoids from *A. reptans* leaves (C+ group, symmetric). We set up 15 replicates per group type. Two hours before the behavioural assay, each individual was colour-marked (green, red, white, or yellow; Uni POSCA marker pens, Japan) on its thorax and wings to enable identification during behaviour scoring. The colours were alternated between sexes and treatments across replicates to avoid any colour effects. In each trial, one to two replicates per group type were set up simultaneously in Petri dishes and recorded for 25 minutes. Agonistic and mating interactions of each individual in a movie replicate were scored by an observer as in experiment 2.

For the social network analysis, we analysed group as well as individual measures of the network. Therefore, we constructed an undirected weighted network for the social interactions (sum of agonistic and mating interactions) in each replicate. We calculated the network weights as the total number of interactions between each pair of individuals (dyad). We normalised the weights at the replicate level and at the dyadic level in each replicate, by dividing by the maximum number of interactions across all replicates and all dyads, respectively. At the group level, we calculated the density of social interactions in each replicate, as following:

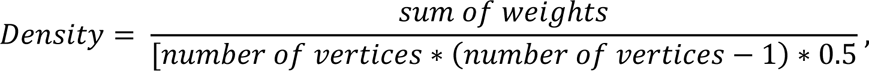

where vertices is the number of individuals in a replicate. Moreover, at the group level, we counted the number of distinct dyads that exhibited social interactions in each replicate. At the individual level, we calculated the strength (Barrat *et al*. 2004) and harmonic centrality (Marchiori & Latora 2000). The strength is a measure of how strongly an individual is directly connected to other individuals in the network. Harmonic centrality calculates the ‘average’ distance of an individual to the other individuals in the network. We used harmonic centrality as we had disconnected networks in some replicates.

### Experiment 4. Costs of direct and indirect clerodanoid acquisition on lifespan, lipid and carbohydrate content

To examine fitness costs of clerodanoid acquisition and effects on metabolism, we measured the lifespan and lipid and carbohydrate content of individuals from different groups. Similar to experiment 3, freshly eclosed *A. rosae* adults were colour-marked and assigned to C- or C+, offering the latter *A. reptans* leaves for 48 hours. The adults were then used to construct three group types consisting of two males and two females, as in experiment 3 (C-group, Mixed group and C+ group). We set up 10 replicates per group type. After four days of being in a group, the adults were separated and one male and one female were taken to score their survivorship while the other male and female were taken to measure their lipid and carbohydrate content. Note that some individuals died while being in a group, they were included for survivorship and metabolic trait analysis.

Lipid and carbohydrate contents were determined for each individual using a method adapted from Cuff *et al*. (2021). Individuals were isolated in Eppendorf tubes (2 ml) and frozen at - 80°C, lyophilised for 48 hours and weighed. Afterwards, each individual was soaked in 0.5 ml of a 1:12 chloroform:methanol solution for 24 hours. From the supernatant 250 µl was collected for lipid analysis, followed by additional soaking of the adult in 0.25 ml of 1:12 chloroform:methanol to remove residual lipids for later carbohydrate analysis. The solvent with residuals was discarded. Lipid analysis was conducted with 50 µl of supernatant in a 96-well plate to which 10 µl of concentrated sulfuric acid was added. The plate was heated in at 100°C for 10 min and then 240 µl of 0.2 mg/ml vanillin in phosphoric acid was added, followed by photometric analysis at 490 nm. A dilution series of cricket oil (Acheta Cricket Oil, Thailand Unique) in 1:12 chloroform:methanol was measured on the same plate for calibration. For carbohydrate analysis, the remaining individuals were homogenised with glass beads, treated with 0.5 ml of 0.1 M NaOH, incubated in a shaker (250 rpm for 30 min at 80 °C) and subsequently allowed to stand at room temperature for 16 hours. Carbohydrate analysis was conducted with 40 µl of the supernatant in a 96-well plate, adding 160 µl of anthrone reagent, and heating the plate at 92°C for 10 min, before photometric analysis at 620 nm. A dilution seried of 1:1 trehalose:glycogen (trehalose: 98%, Roth; glycogen: AppliChem) in water was measured on the same plate for calibration of carbohydrates. Body lipid and carbohydrate content were determined as percentage relative to dry body mass.

We also conducted a similar experiment constructing pairs of females with no access, asymmetric access or symmetric access to clerodanoids (see SI.2), which allowed us to confirm that our effects of direct and indirect clerodanoid acquisition on lifespan (see Results) were not stemming only from increased mating interactions in Mixed and C+ groups but also from increase agonistic interactions in these group types.

### Statistical analyses

All statistical analyses were conducted using R version 4.2.1 (R Core Team 2022). We used Poisson-distributed generalised linear mixed models (GLMM) with trial as a random effect to assess the effect of female treatment on agonistic and mating interactions in Experiment 2 (package ‘lme4’ version 1.1-30, package ‘lmerTest’ version 3.1-3; Bates *et al*. 2015; Kuznetsova *et al*. 2017). Effect of female treatment on mating latency was examined using a linear mixed model (LMM) with trial as a random effect. Differences in clerodanoid 1 and 2 among treatment groups were assessed using Kruskal-Wallis tests followed by Dunn tests with Holm correction (‘FSA’ package version 0.9.4, Ogle *et al*. 2023). In Experiment 3, we ran LMMs for density and binomial GLMMs for the number of dyads with social interactions, with group type as a fixed effect and trial as a random effect. Effect of group type, sex and their interaction on individual-level network measures were analyzed separately for C- and C+ sawflies using poisson-distributed GLMMs for strength and LMMs for harmonic centrality (‘igraph’ package version 1.5.1, Csárdi *et al*. 2023), with replicate ID and trial as random effect. For Experiment 4, individuals were classified based on both their individual and group treatments, yielding four composite treatment categories: “C-from C- group”, “C-from Mixed group”, “C+ from Mixed group”, and “C+ from C+ group”. LMM were used to assess the impact of sex, composite treatment, and their interaction on lifespan, lipid, and carbohydrate content, with replicate ID as random effect. Model validity was confirmed by examining residual distributions. If the interaction terms were non-significant, they were dropped to test the significance of the lower-order terms. Post-hoc analyses were conducted using the ‘multcomp’ package version 1.4-20 (Hothorn *et al*. 2008) for LMMs and GLMMs.

## Results

### Intraspecific variation in clerodanoids in natural A. rosae populations

Both females and males showed intraspecific variation in the amount of putative clerodanoid 1 and clerodanoid 2 across the three locations from which they were collected (Figure 1). Some individuals had no clerodanoids, while others had relatively high quantities.

**Figure 1.**
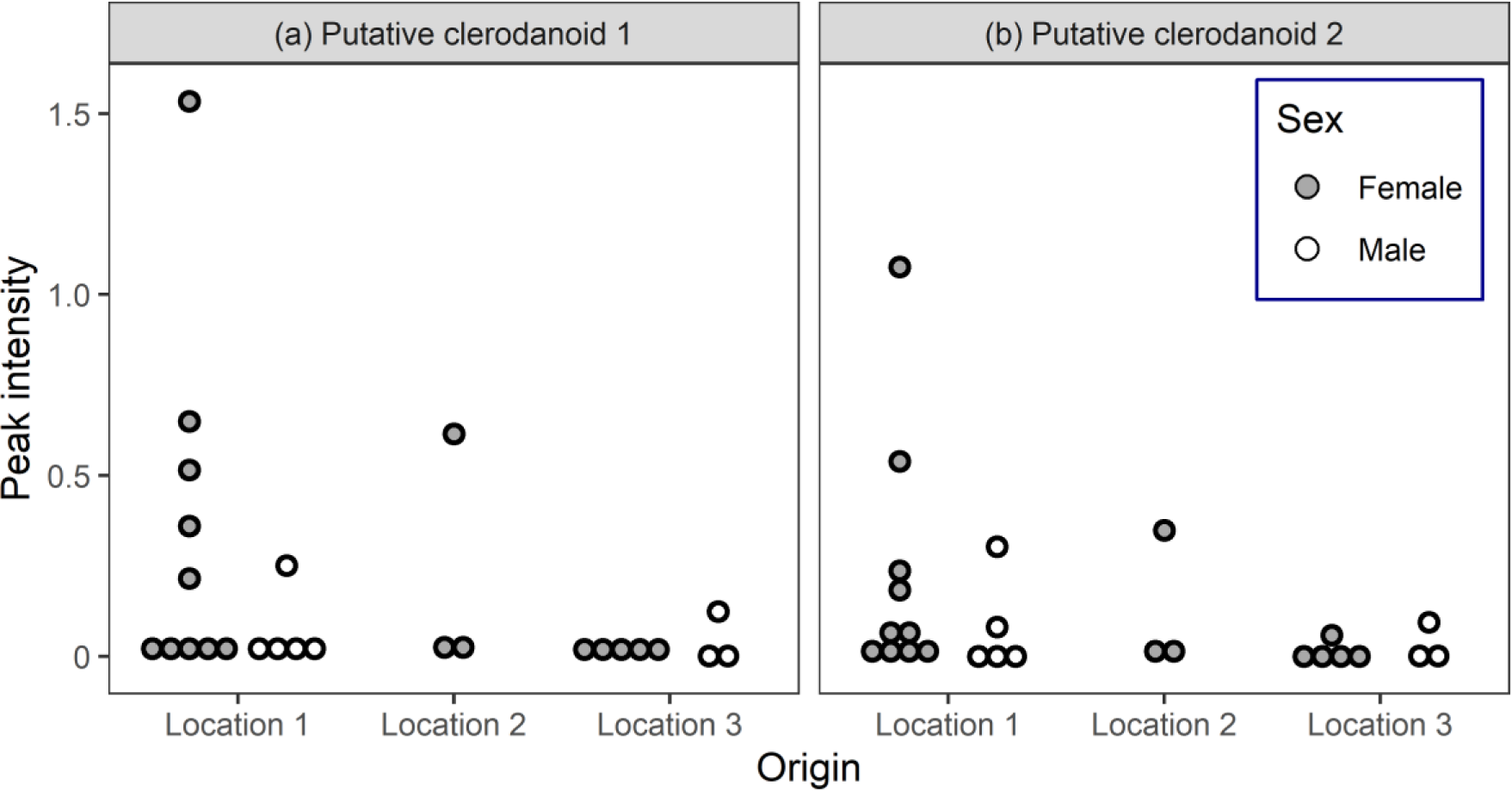
Intraspecific variation in peak intensities of putative (a) clerodanoid 1 and (b) clerodanoid 2 per individual in wild-caught female (n=18) and male (n=8) *Athalia rosae* sawflies from three locations.

### Clerodanoid uptake affects sawfly behaviour and varied levels observed in lab-reared sawflies

There was a significant effect of female treatment on number of agonistic interaction occurrences (*χ^2^* = 48.56, df = 2, *P* < 0.001, Figure 2a), but not on mating occurrences (*χ^2^* = 1.76, df = 2, *P* = 0.41, Figure 2b). C- and ‘C+ from conspecific’ treatment females had significantly fewer agonistic interactions than ‘C+ from plant’ treatment females (SI.3a). There was a significant effect of female treatment on mating latency (*χ^2^*= 9.90, df = 2, *P* = 0.007, Figure 2c) with C- females having a significantly longer latency until mating than ‘C+ from plant’ females, while ‘C+ from conspecific’ females showed an intermediate mating latency (SI.3b).

**Figure 2.**
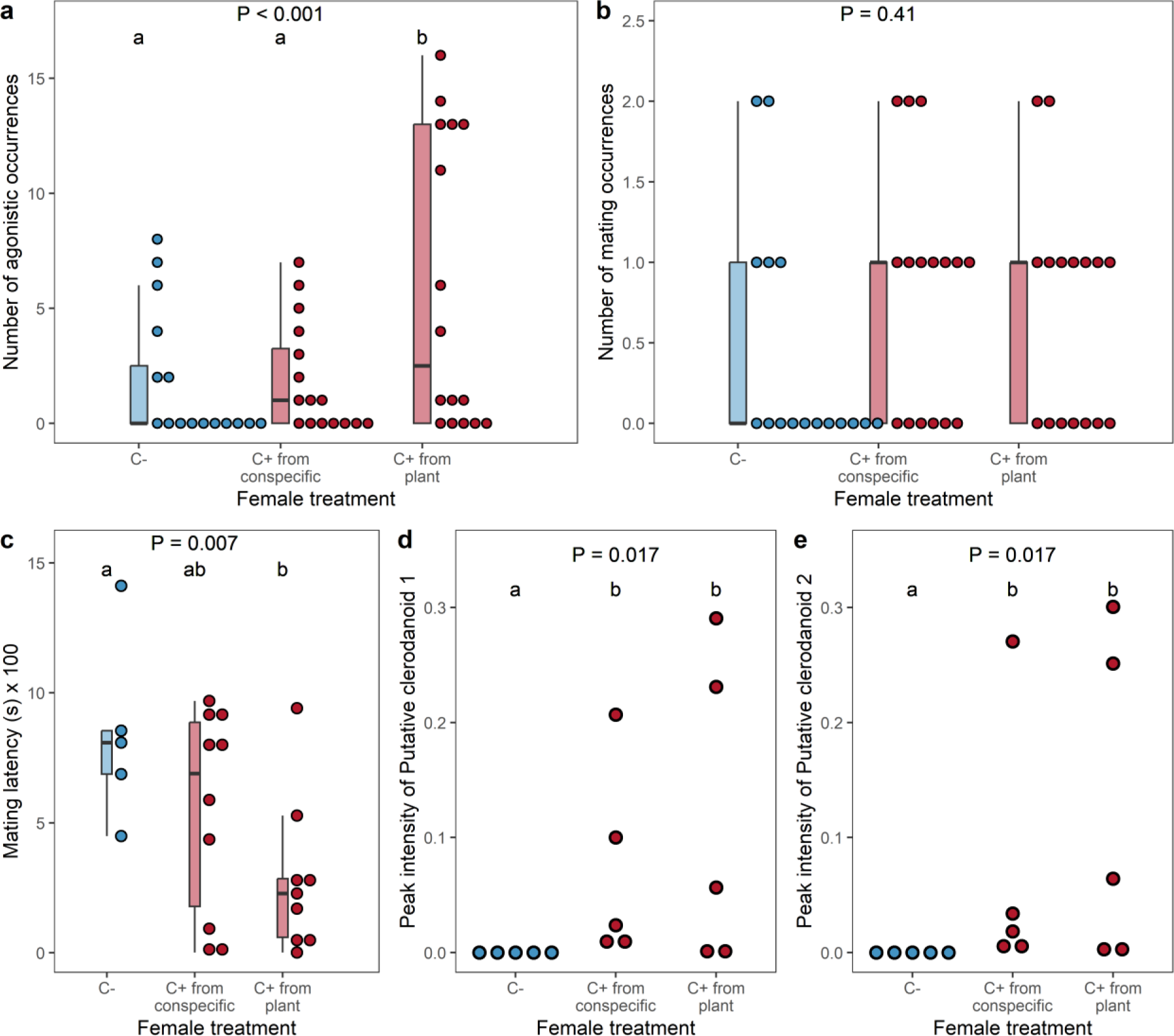
Effect of female treatment on the number of occurrences of (a) agonistic and (b) mating interactions, as well as (c) mating latency with C- males of *Athalia rosae*. Additionally, the effect of female treatment on the peak intensity of putative (d) clerodanoid 1 and (e) clerodanoid 2 in lab-reared female sawflies is shown. Treatments include no access (C-), indirect access (C+ from conspecific: prior contact with other females that had access to *Ajuga reptans* leaf), or direct access (C+ from plant: prior access to *A. reptans* leaf) to clerodanoids. Boxplots display the median, 25th, and 75th percentiles, with whiskers indicating values within 1.5 times the interquartile range. Coloured circles represent individual data points. *P*-values are derived from GLMM (a,b), LMM (c), or Kruskal-Wallis (d,e) tests. Different lowercase letters indicate significant differences from post-hoc tests (*P* < 0.05).

There was a significant difference between C-, ‘C+ from conspecific’ and ‘C+ from plant’ females in the amount of clerodanoid 1 (*χ^2^*= 8.11, df = 2, *P* = 0.017, Figure 2d) and clerodanoid 2 (*χ^2^*= 8.10, df = 2, *P* = 0.017, Figure 2e). No clerodanoids were detected in C-females, while there was intraspecific variation in clerodanoids in both ‘C+ from conspecific’ and ‘C+ from plant’ females, which did not differ significantly from each other (SI.3c, SI.3d).

### Intraspecific variation in clerodanoid access modulates social networks

The group types varied in their social networks, with the C- group having fewer interactions than the Mixed and the C+ group (Figure 3a). Our results showed that there was a significant effect of group type on density (*χ^2^* = 27.98, df = 2, *P* < 0.001, Figure 3b) and number of dyads with social interactions (*χ^2^* = 41.29, df = 2, *P* < 0.001, Figure 3c). Mixed groups had a significantly higher density and more dyads that had interacted than C- and C+ groups (SI.4a, SI.4b). Together this suggests that access to clerodanoids increases social interactions with more individuals networking, particularly in the Mixed group.

**Figure 3.**
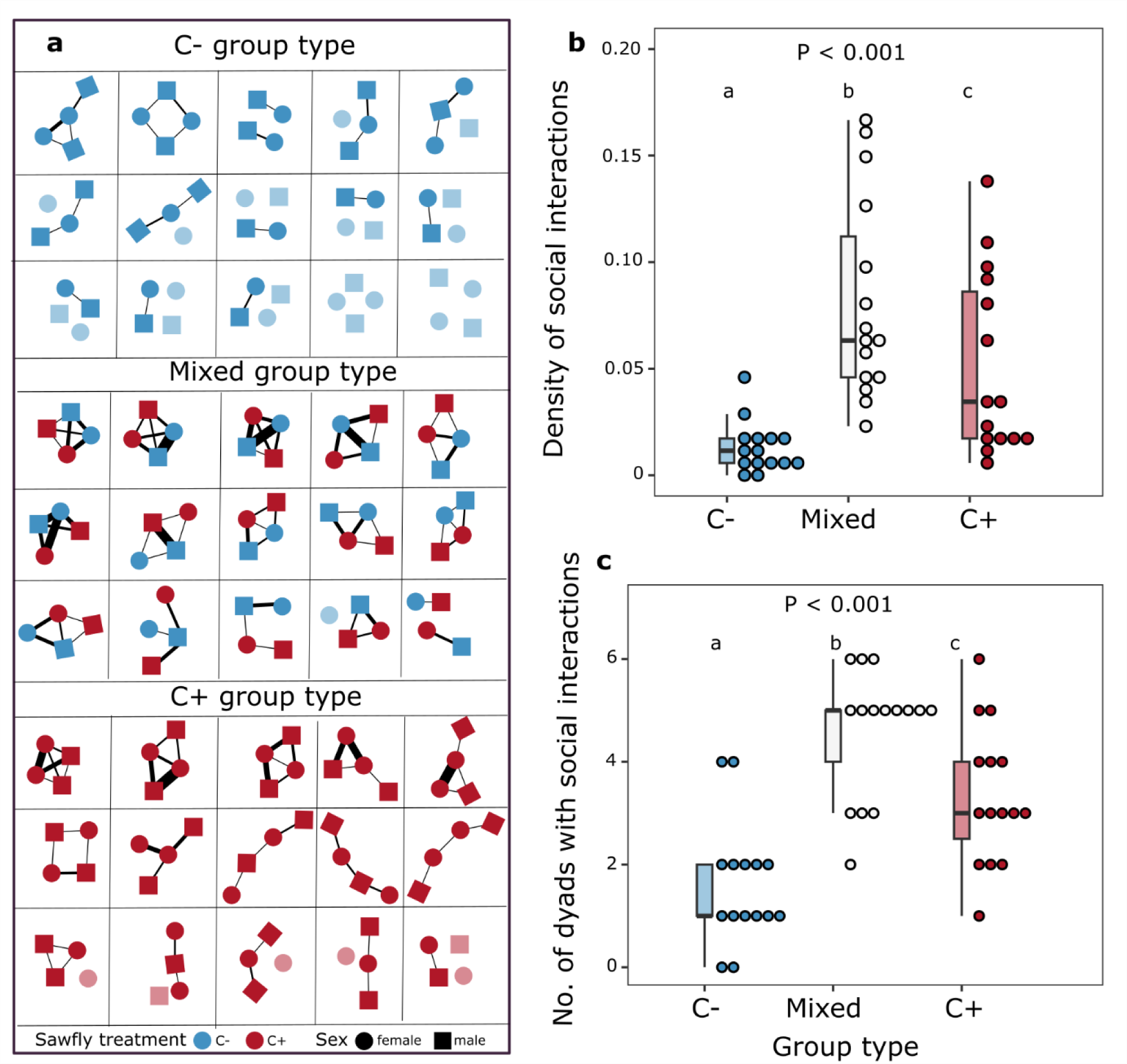
(a) Network representations of C-, Mixed and C+ groups (n=15 per group type). Each cell in the grid corresponds to one replicate and consisted of two female (circular nodes) and two male (square nodes) adults of *Athalia rosae*. Sawflies had either no access (C-, red nodes) or prior access to *Ajuga reptans* (C+, blue nodes). Social interactions between dyads are depicted by lines, with line thickness indicating interaction frequency. Lighter nodes indicate individuals with no social interactions during the observation period. Effect of group type on (b) density of social interactions and (c) number of dyads with social interactions. Boxplots display the median, 25th, and 75th percentiles, with whiskers indicating values within 1.5 times the interquartile range. Coloured circles represent individual data points. *P*-values are derived from LMM (b) or GLMM (c) models. Different lowercase letters denote significant differences from post-hoc tests (*P* < 0.05).

For C- sawflies, there was a significant effect of group type (*χ^2^* = 63.55, df = 1, *P* < 0.001) on strength (Figure 4a) such that C- sawflies in Mixed groups had a higher strength, i.e., had more social interactions, than adults in C- groups (SI.5a). In contrast, there was a significant effect of sex (*χ^2^* = 25.14, df = 1, *P* < 0.001) on strength of interactions (Figure 4b) for C+ sawflies, with males having a lower strength than females across Mixed and C+ groups (SI.5a). For harmonic centrality, there was a significant effect of group type (*χ^2^* = 6.88, df = 1, *P* = 0.008) in C- sawflies (Figure 4c), such that those in Mixed groups had higher harmonic centrality values. There was no significant effect of any predictor variable on harmonic centrality in C+ sawflies (Figure 4d, SI.5c).

**Figure 4.**
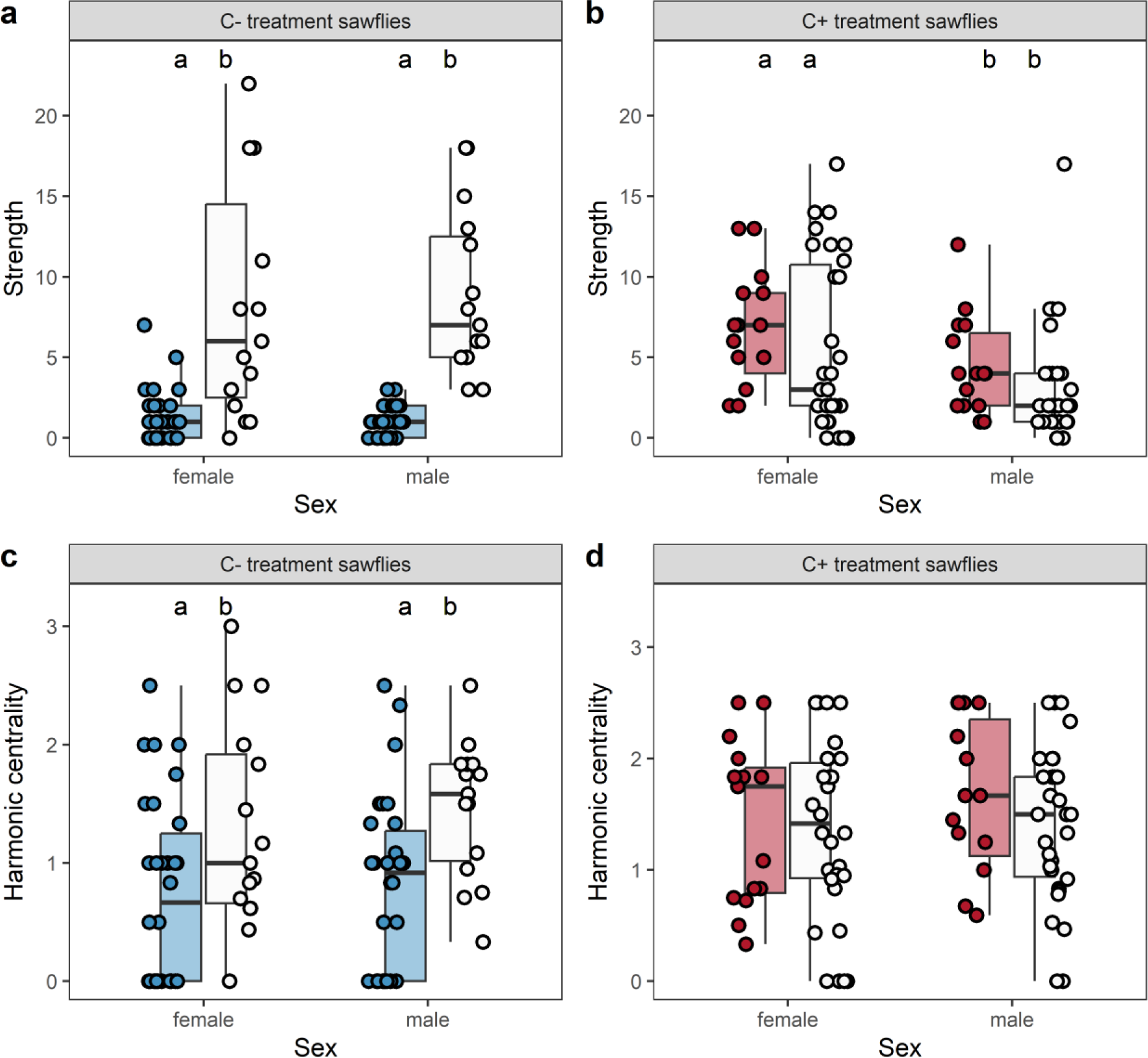
Effect of group type (C-, Mixed, or C+ groups, distinguished by different colours) on strength (top row) and harmonic centrality (bottom row) in C- treatment (a, c) and C+ treatment (b, d) *Athalia rosae* sawflies of both sexes. Boxplots depict the median, 25th, and 75th percentiles, with whiskers indicating values within 1.5 times the interquartile range. Coloured circles denote individual data points. Different lowercase letters indicate significant differences from post-hoc tests (*P* < 0.05).

### C+ from Mixed group adults have a shorter lifespan but no effect on lipid and carbohydrate content

For lifespan, there was a significant effect of composite treatment (*χ^2^* = 18.74, df = 3, *P* < 0.001) such that “C+ from Mixed group” sawflies had a shorter lifespan than “C-from C- group”, “C-from Mixed group”, and “C+ from C+ group” (Figure 5a, SI.6a,7). Additionally, sex (*χ^2^*= 12.52, df = 1, *P* < 0.001) had a significant effect on lifespan with males having a shorter lifespan than females. There was no significant effect of sex, composite treatment or their interaction on body lipid content (SI.6b). Only sex had a significant effect on body carbohydrate content (*χ^2^*= 13.09, df = 1, *P* < 0.001; SI.6c), with females having higher carbohydrate content than males. We obtained qualitatively similar results for the effect of direct and indirect clerodanoid acquisition on lifespan using female pairs (see SI.2). Together, this suggests that social interactions, especially agonistic, to obtain clerodanoids may have fitness costs in terms of lifespan.

**Figure 5.**
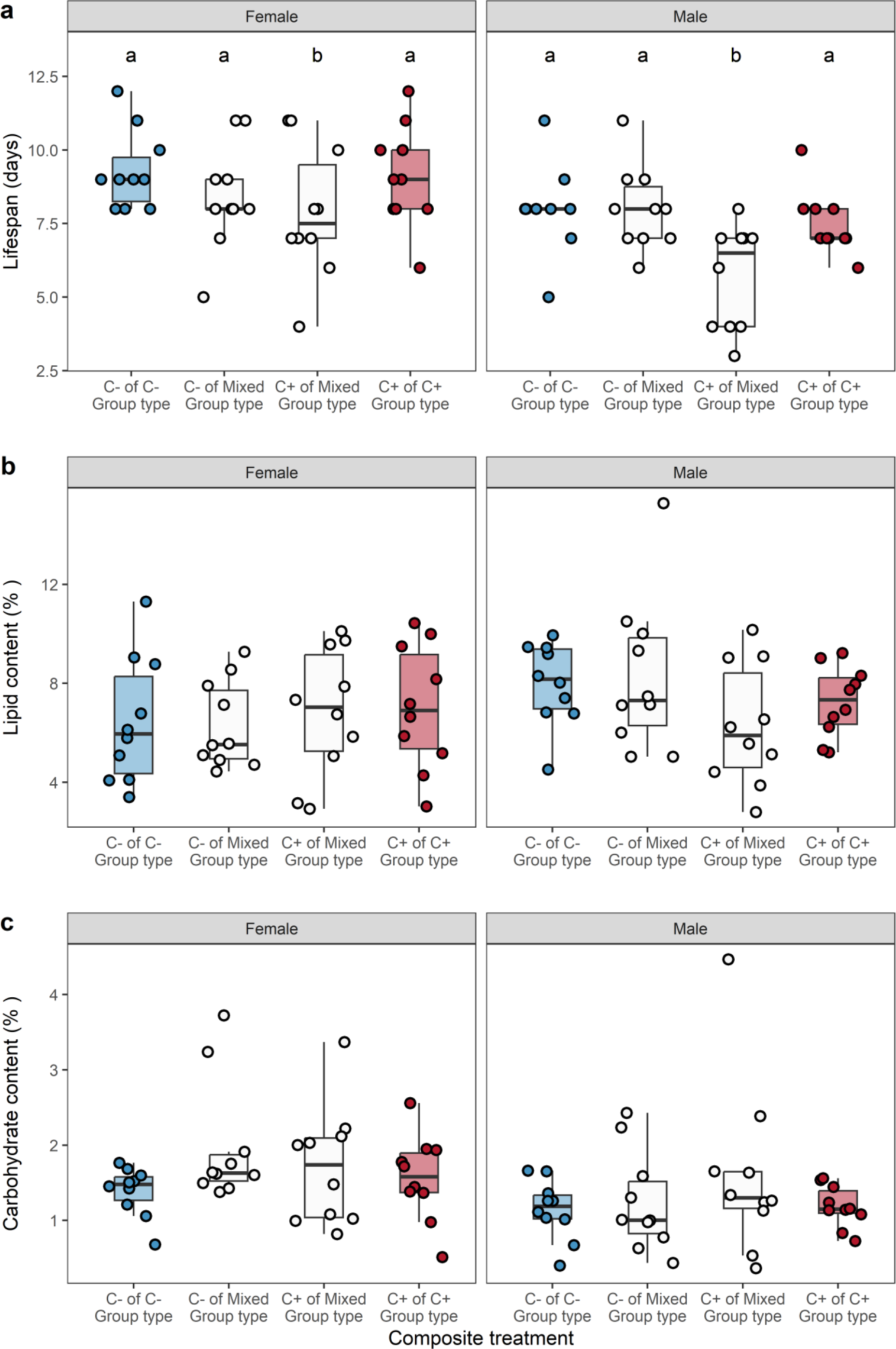
Effect of composite treatment on (a) lifespan, (b) carbohydrate content (%), and (c) lipid content (%) in female and male *Athalia rosae* sawflies. Sawflies were assigned to either C- or C+ treatment groups (C-: no access, C+: prior access to *Ajuga reptans* leaf) and grouped as C-, Mixed, or C+ groups. Boxplots display the median, 25th and 75th percentiles, with whiskers indicating values within 1.5 times the interquartile range. Coloured circles denote individual data points. Different lowercase letters represent significant differences from post-hoc tests (*P* < 0.05).

## Discussion

We observed variation in clerodanoid amounts in wild-caught *A. rosae* adults of both sexes and from all locations. A capture release study had shown that most *A. rosae* adults could be recaptured from within 100 m of their release, suggesting limited dispersal in this species (Nagasaka 1992). Thus, our field populations were likely separated, indicating that intraspecific variation in sequestered compounds may be common in natural populations of herbivorous insects that are able to sequester plant metabolites. Moreover, we observed intraspecific variation in clerodanoid amounts of lab-reared sawflies with both direct (via plant leaves) and indirect (via nibbling on conspecifics) access to clerodanoids, similar to previous studies (Singh *et al*. 2022). Such intraspecific variation in sequestered plant chemicals in animals is widely observed (Opitz & Müller 2009; Speed *et al*. 2012). While this variation may result from environmental stochasticity, e.g., variation in metabolite concentrations across plants, it could also have an evolutionarily basis, e.g., some individuals of an animal species may be chemically defended while others are not (Brower *et al*. 1970; Speed *et al*. 2012). Our study shows that intraspecific variation in sequestered plant chemicals can impact social networks and have fitness costs.

The presence and quantity of clerodanoids can be considered as important parts of the chemical phenotype of individuals, which could play a key role in shaping the individualised niche, as has been argued for plants (Müller & Junker 2022). While effects of food plants on the phenotype and behaviour of herbivorous individuals are widely documented (Geiselhardt *et al*. 2012; Jarrett & Miller 2024; Müller & Müller 2017; Singh *et al*. 2020), we showed here that plant chemicals sequestered from plants that do not serve as food plants can also modulate social interactions. Our results demonstrated that social networks comprising Mixed groups exhibited the highest density and number of interacting dyads, followed by C+ groups. The asymmetric access to clerodanoids in Mixed groups may lead to an increase in social interactions, potentially due to C- trying to obtain clerodanoids from C+ individuals. Moreover, C- individuals in Mixed groups could acquire clerodanoids from C+ individuals, which may then increase their social interactions, as it will change their clerodanoid acquisition status. Individuals in C- groups had the least density and number of interacting dyads, suggesting that when all individuals lack clerodanoids, they have only few intra-and inter-sex social interactions.

The social interactions of the individuals depended not only on their own clerodanoid acquisition status but also on that of other conspecifics in the group. For example, C- individuals had a higher number of social interactions in Mixed than in C- groups (see Figure 4a,4c), suggesting that even for the same clerodanoid-holding status, an individual may have low or high frequencies of social interactions, depending on the group composition. Similar effects of conspecifics on an individual’s behaviour have been shown in an aquatic insect, *Notonecta irrorata*, where the propensity to disperse depended on the behavioural type of conspecifics present (Kitchen & Chalcraft 2020). Indeed, studies across various animal species have shown that the behaviour of a focal individual can be shaped by both internal factors, such as personality traits, and external factors, such as social context and interactions with conspecifics (Webster & Ward 2011). Given the intraspecific variation in clerodanoid content that we observed in wild-caught individuals of *A. rosae*, we expect social interactions in natural populations to range from low, for example, if no *A. reptans* plants are around from which clerodanoids can be sequestered, to high, when some insects could sequester clerodanoids, leading to a skewed distribution of social interaction frequency.

Social interactions may accrue over time, and influence fitness of individuals through different ways, such as injury or mortality resulting from agonistic interactions, variation in mating frequency or disease progression (Alwash & Levine 2019; Dawson *et al*. 2018; Guo & Dukas 2020; Stroeymeyt *et al*. 2018). We observed a reduced lifespan in C+ adults from Mixed groups. In experiment 2, we saw that C+ females paired with C- males had the most agonistic interactions. Similarly, in Mixed groups, we found more frequent interactions than in C- or C+ groups. Thus, we can expect that C+ adults in Mixed groups had more social, especially agonistic, interactions with their conspecifics that may have nibbled on the C+ individual to get access to clerodanoids. These increased agonistic interactions may have led to a decreased lifespan for C+ adults in Mixed groups. Such costs of aggression have also been shown in the fruit fly, *Drosophila melanogaster*, where flies that were engaged in aggressive behaviour had a reduced survival despite not showing any differences in wing damage from no-aggression treatment flies (Guo & Dukas 2020). This particular study hypothesised that such costs could arise from physiological and metabolic changes triggered by stress and aggressive conflicts. Indeed, a previous study (Paul *et al*. 2021) demonstrated transcriptional upregulation of metabolic pathway genes in *A. rosae* sawflies engaging in conspecific nibbling, indicating potential costs of aggressive interactions. However, we found no significant impact on body lipid and carbohydrate content in the present study.

Along with chemical defence against predation and increased mating success, sequestered plant metabolites may also lead to benefits for the offspring (Eisner *et al*. 2000; Eisner & Meinwald 1995; Gonzalez *et al*. 1999). However, at least under controlled laboratory conditions, no significant effects of parental clerodanoid status on offspring was found in *A. rosae* (Paul *et al*. 2022). Thus, while clerodanoid acquisition may be associated with benefits such as enhanced defence against predation (Singh *et al*. 2022) and mating success (Amano *et al*. 1999; Paul & Müller 2021), there may be a trade-off with costs in terms of a reduced lifespan due to agonistic interactions. This in turn may maintain intraspecific variation in clerodanoids in natural populations of *A. rosae*. Understanding the effects of variation in these sequestered plant metabolites is essential because these ecological effects can scale up from the individual level to factors that shape animal social interaction networks.

## Acknowledgements

This study was funded by the German Research Foundation (DFG) as part of the SFB TRR 212 (NC^3^), project number 396777467 (granted to CM). We thank Karin Djendouci for her help in conducting the experiments and Rohit Sasidharan for help with collecting sawflies.

## Author Contributions

Conceptualization and experimental design: PS; data collection: YS for experiment 2, SJ for experiment 3, SJ, PS and LB for experiment 4; chemical analysis: LB; data validation and analysis: PS and GB for network analysis, PS for other experiments; data visualisation: PS; writing original draft: PS, CM; reviewing and editing: PS, CM, LB, GB; funding acquisition: CM.

## Data availability

The datasets are available on the Zenodo repository (https://doi.org/10.5281/zenodo.10969832).

