## Supplementary information for "Plant metabolites modulate animal social networks and lifespan"

**SI.1.** Locations of wild-caught *Athalia rosae* adults.

| Location | Sex | Number of individuals collected |
| --- | --- | --- |
| Location 1: around 52°02'40.6"N 8°29'32.3"E | Female | 10 |
| Location 1: around 52°02'40.6"N 8°29'32.3"E | Male | 5 |
| Location 2: around 52°02'51.2"N 8°30'55.4"E | Female | 3 |
| Location 3: around 52°03'36.0"N 8°34'55.2"E | Female | 5 |
| Location 3: around 52°03'36.0"N 8°34'55.2"E | Male | 3 |

**SI.2.** We conducted an experiment to confirm the costs of direct and indirect clerodanoid acquisition on lifespan in female *A. rosae* sawflies. Using only female *A. rosae* sawflies allowed us to examine the costs of clerodanoid acquisition independent of mating effects.

**Methods**

We collected freshly eclosed females and assigned them to either C- or C+ treatment for 48 hours. Next, we put together either two C- individuals (C-C- pair, 28 replicates), a C- and a C+ individual (C-C+ pair, 30 replicates) or two C+ individuals (C+C+ pair, 29 replicates). The C- individual from the C-C+ pair could acquire clerodanoids from the C+ conspecific. Individuals in a replicate were marked with green or white colour paint to allow for differentiation. After 48 hours, we randomly chose one individual from each replicate of the C-C- pair and the C+C+ pair, and both individuals from the C-C+ pair. Each individual was placed in a Petri dish and checked daily for survivorship.

**Statistical analyses**

To analyse differences in lifespan of C- females (from C-C- pair), C- and C+ females (from C-C+ pair) and C+ females (from C+C+ pair), we produced Kaplan-Meier plots and compared the curves by the log-rank test using ‘survival’ package version 3.5-0 (Therneau 2023) and ‘survminer’ package version 0.4.9 (Kassambara *et al.* 2021). Similarly, we tested for differences in lifespan between C+ individuals from C+C+ pairs and C+ individuals from C-C+ pairs.

**Results**

***C+ individuals from C-C+ pairs have a shorter lifespan***

There was no significant difference between the lifespan of C- females from C-C- pairs, C- females from C-C+ pairs and C+ females from C+C+ pairs (*χ_2_^2^* = 2.3, *P* = 0.3, Figure 5). However, C+ adults from C-C+ pairs had a significantly shorter lifespan than C+ adults from C+C+ pairs (*χ_1_^2^* = 5.2, *P* = 0.02), suggesting that agonistic interactions to obtain clerodanoid may have fitness costs.

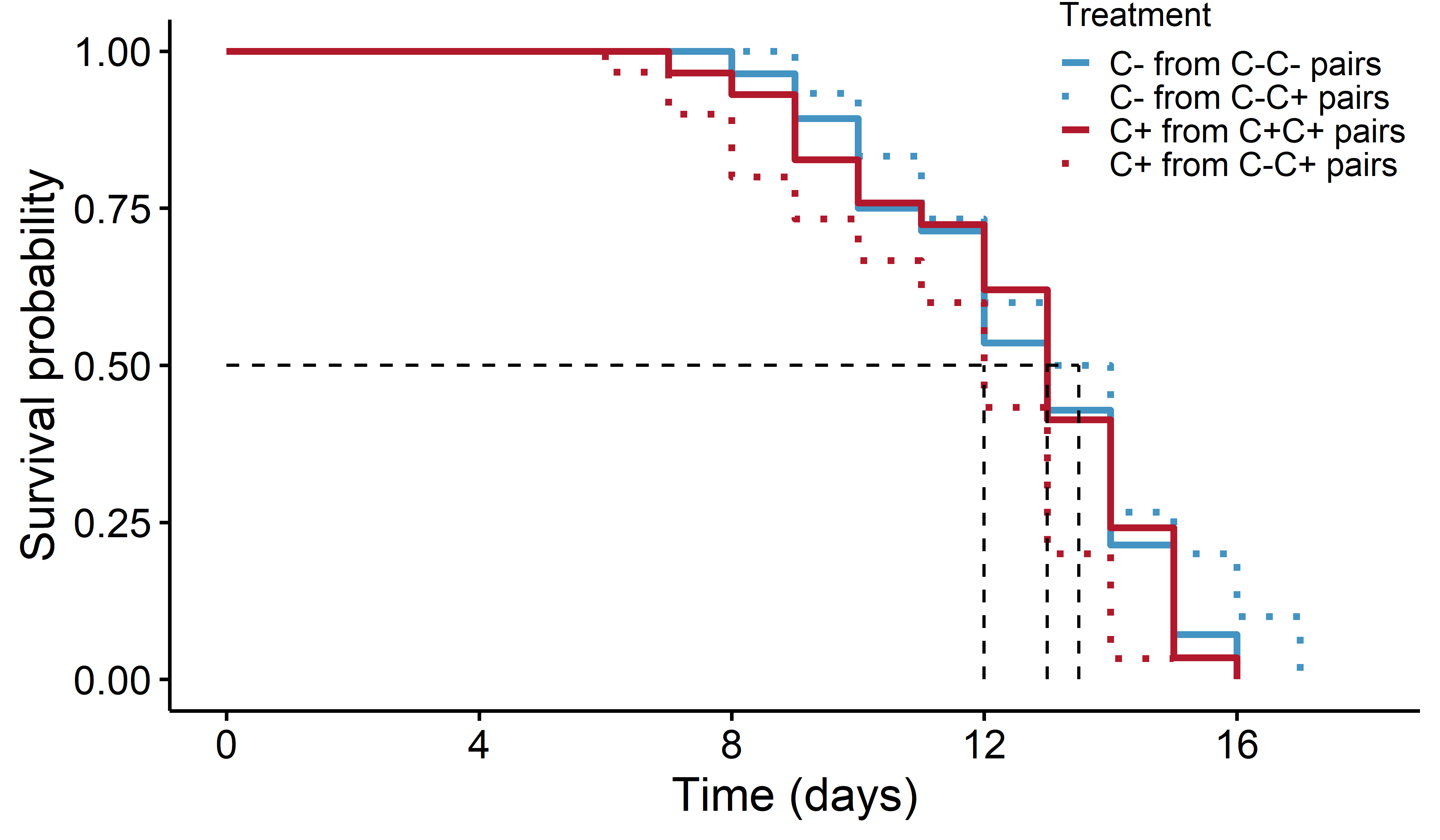

Figure SI.2.1. Survival of *Athalia rosae* females kept for 48 h with an individual of the same or a different treatment (C-: no access, C+: prior access to *Ajuga reptans* leaf), represented by Kaplan-Meier curves. The curve depicts the proportion of females surviving over time. The black dashed line represents the median survival of each treatment given by the point where it intersects the treatment curve. Note that the C- individuals from C-C+ pairs and C+ individuals from C-C+ pairs originate from the same treatment (see above text for details).

**SI.3.** Results of post-hoc analyses from examining the effect of female treatment on the (a) number of agonistic interactions, (b) mating latency, (c) (putative) clerodanoid 1 and (d) clerodanoid 2 in experiment 2. *P*-values adjusted with the Holm method. Significant differences (*P* < 0.05) are highlighted in bold.

| Variable | Pairwise comparison | *z* value | *P* |
| --- | --- | --- | --- |
| a) Number of occurrences of agonistic interactions | AC+ - C- | 0.13 | 0.895 |
|  | C+ - C- | 5.53 | **< 0.001** |
|  | C+ - AC+ | 5.44 | **< 0.001** |
| b) Mating latency | AC+ - C- | -1.52 | 0.128 |
|  | C+ - C- | -3.08 | **0.006** |
|  | C+ - AC+ | -1.91 | 0.110 |
| c) Clerodanoid 1 | AC+ - C- | 2.51 | **0.035** |
|  | C+ - C- | 2.40 | **0.031** |
|  | C+ - AC+ | -0.10 | 0.912 |
| d) Clerodanoid 2 | AC+ - C- | 2.44 | **0.028** |
|  | C+ - C- | 2.48 | **0.039** |
|  | C+ - AC+ | 0.03 | 0.970 |

**SI.4.** Result of post-hoc analyses for examining the effects of group type on (a) density and (b) number of dyads/pairs with social interactions in social networks in experiment 3. *P*-values adjusted with the Holm method. Significant differences (*P* < 0.05) are highlighted in bold.

| Variable | Pairwise comparison | *z* value | *P* |
| --- | --- | --- | --- |
| a) Density | C+ - C- | 2.89 | **0.007** |
|  | Mix - C- | 5.28 | **< 0.001** |
|  | C+ - Mix | -2.39 | **0.016** |
| b) Number of dyads with social interactions | C+ - C- | 3.90 | **< 0.001** |
|  | Mix - C- | 6.39 | **< 0.001** |
|  | C+ - Mix | -2.98 | **0.002** |

**SI.5.** Effects of predictor variables on individual social network parameters of (a) strength and (b) harmonic centrality in sawfly treatment of C- and C+. Significant effects (*P* < 0.05) are highlighted in bold. We dropped non-significant interaction terms to test the significance of the lower-order terms.

| Network parameter | Sawfly treatment | Predictor variables | *X^2^* | *df* | *P* |
| --- | --- | --- | --- | --- | --- |
| a) Strength | C- | Group type * sex | 1.23 | 1 | 0.266 |
|  |  | Group type | 63.55 | 1 | **< 0.001** |
|  |  | Sex | 0.00 | 1 | 1 |
|  | C+ | Group type * sex | 0.44 | 1 | 0.506 |
|  |  | Group type | 2.90 | 1 | 0.088 |
|  |  | Sex | 25.14 | 1 | **< 0.001** |
| b) Harmonic centrality | C- | Group type * sex | 0.23 | 1 | 0.629 |
|  |  | Group type | 6.88 | 1 | **0.008** |
|  |  | Sex | 0.55 | 1 | 0.457 |
|  | C+ | Group type * sex | 0.56 | 1 | 0.451 |
|  |  | Group type | 0.52 | 1 | 0.469 |
|  |  | Sex | 0.87 | 1 | 0.349 |

**SI.6** Effects of predictor variables on lifespan, lipid content (%) and carbohydrate content (%) in *A. rosae* adults. Significant effects (*P* < 0.05) are highlighted in bold. We dropped non-significant interaction terms to test the significance of the lower-order terms.

|  | Predictor variables | *X^2^* | *df* | *P* |
| --- | --- | --- | --- | --- |
| (a) Lifespan | Group type * sex | 4.36 | 3 | 0.224 |
|  | Group type | 18.74 | 3 | **< 0.001** |
|  | Sex | 12.52 | 1 | **< 0.001** |
| (b) Lipid content (%) | Group type * sex | 3.43 | 3 | 0.329 |
|  | Group type | 1.84 | 3 | 0.606 |
|  | Sex | 2.79 | 1 | 0.094 |
| (c) Carbohydrate content (%) | Group type * sex | 3.80 | 3 | 0.283 |
|  | Group type | 1.54 | 3 | 0.672 |
|  | Sex | 13.09 | 1 | **< 0.001** |

**SI.7.** Result of post-hoc analyses for examining the effects of treatment on lifespan in *A. rosae* adults. *P*-values adjusted with the Holm method. Significant differences (*P* < 0.05) are highlighted in bold.

| Variable | Pairwise comparison | *z* value | *P* |
| --- | --- | --- | --- |
| Lifespan | C- from Mixed Group type - C- from C- Group type | -0.776 | 0.656 |
|  | C+ from C+ Group type - C- from C- Group type | -0.601 | 0.657 |
|  | C+ from Mixed Group type - C- from C- Group type | -3.931 | **< 0.001** |
|  | C+ from C+ Group type - C- from Mixed Group type | 0.175 | 0.861 |
|  | C+ from Mixed Group type - C- from Mixed Group type | -3.155 | **0.003** |
|  | C+ from Mixed Group type - C+ from C+ Group type | -3.329 | **0.002** |
